# Can the microbiome influence host evolutionary trajectories?

**DOI:** 10.1101/700237

**Authors:** Lucas P. Henry, Marjolein Bruijning, Simon K.G. Forsberg, Julien F. Ayroles

## Abstract

The microbiome shapes many traits in hosts, but we still do not understand how it influences host evolution. To impact host evolution, the microbiome must be heritable and have phenotypic effects on the host. However, the complex inheritance and context-dependence of the microbiome challenges traditional models of organismal evolution. Here, we take a multifaceted approach to identify conditions in which the microbiome influences host evolutionary trajectories. We explore quantitative genetic models to highlight how microbial inheritance and phenotypic effects can modulate host evolutionary responses to selection. We synthesize the literature across diverse taxa to find common scenarios of microbiome driven host evolution. First, hosts may leverage locally adapted microbes, increasing survivorship in stressful environments. Second, microbial variation may increase host phenotypic variation, enabling exploration of novel fitness landscapes. We further illustrate these effects by performing a meta-analysis of artificial selection in Drosophila, finding that bacterial diversity also frequently responds to host selection. We conclude by outlining key avenues of research and experimental procedures to improve our understanding of the complex interplay between hosts and microbiomes. By synthesizing perspectives through multiple conceptual and analytical approaches, we show how microbiomes can influence the evolutionary trajectories of hosts.

## Introduction

The microbiome has emerged as a key determinant of many aspects of organismal biology, capable of shaping developmental, physiological, and reproductive phenotypes^1–5^. Yet, the contribution of the microbiome to host adaptation remains an evolutionary puzzle^6–9^. Microbiomes are traditionally viewed as non-genetic, environmental factors that influence host phenotypes. However, unlike abiotic environmental conditions, the effects of microbial variation have a genetic basis and can evolve^10^, but are not inherited in the same way as host genes^6,7,9^. Frequently, microbiomes have substantial phenotypic effects on their hosts, but these effects strongly depend on the ecological context^2,8,11^. Despite their clear importance, the complex effects of microbial inheritance and genetics on host phenotypes remain underappreciated, limiting our ability to understand host-microbiome evolution.

In this perspective piece, we explore how the microbiome influences host evolutionary trajectories. To assess the importance of the microbiome on host phenotypes and inheritance, we integrate the microbiome into quantitative genetic models to make predictions about the microbial impact on host evolutionary trajectories. We review techniques to measure host-microbiome evolution, highlighting current technical and theoretical limitations. Next, we detail two common scenarios found in the literature where the microbiome influences host evolution across a diverse range of host organisms. First, hosts may leverage locally adapted microbes. Second, microbial variation may increase host phenotypic variation, enabling exploration of novel fitness landscapes. Then, to better understand how microbiomes may influence host evolution, we perform a meta-analysis of experimental evolution in *Drosophila* to demonstrate that microbial diversity frequently responds to host adaptation. We conclude by suggesting key avenues of research and outline experimental approaches to improve our understanding of how and when the microbiome may influence host evolution. Through this perspective piece, we show how understanding host-microbial variation can provide fundamental insights into ecological and evolutionary trajectories.

## Extending the Host Genetic Repertoire

Dawkins’ *Extended Phenotype* recognized how organisms modify surrounding environments and the ecological community^12^. Through environmental modification, an organism’s phenotypic effects are extended beyond its own genome, suggesting evolution is influenced through interacting ecological communities. This theory, developed for free-living ecosystems, also applies to host-microbiome interactions^13,14^. The microbiome, with its consortium of genomes, extends the genetic repertoire of the host to form what some are now calling the *Extended Genotype* because the host integrates the extended effects of the microbiome into its phenotype^11,15–23^ (Fig. 1). The evolutionary impact of the microbiome will depend on how host and microbial genetic variation are inherited and interact, shaping host phenotypes across generations.

**FIGURE 1:**
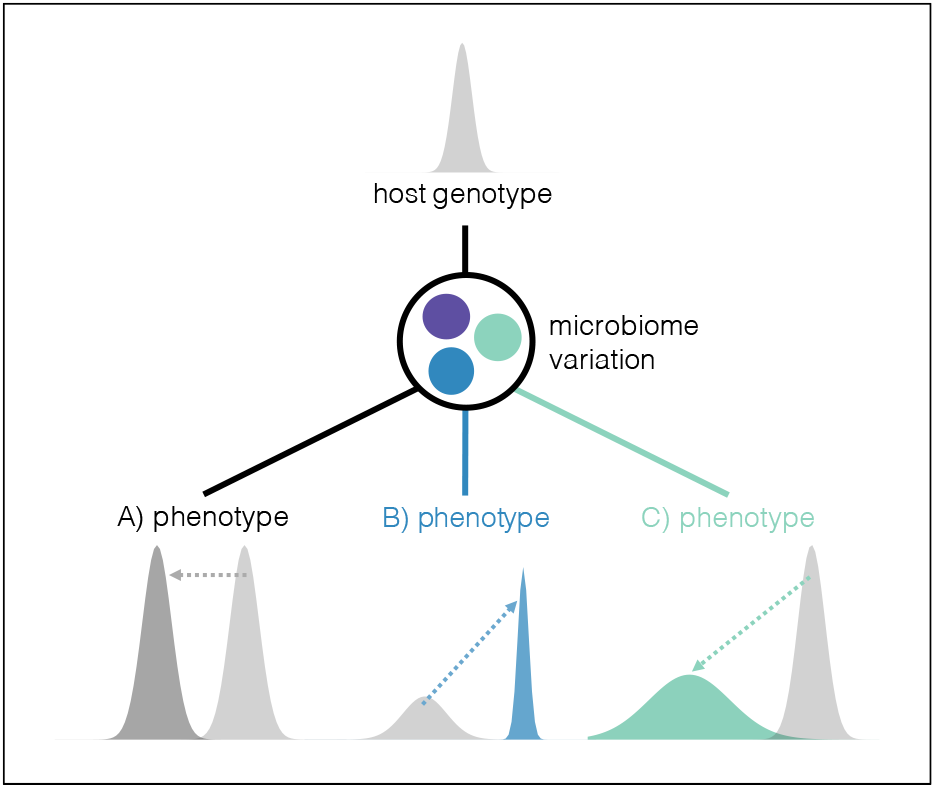
Conceptualization of the microbiome influence on host phenotype. The microbiome encodes many more genes than the host genome alone. Interactions with variation in the microbiome may alter the host genotype-phenotype map, shaping host phenotypic variation. **A)** First, we conceptualize the phenotypic variation encoded by the host genome alone in light grey. The presence of a microbiome might shift the mean of host phenotypes, visualized as the by the arrow to the darker grey distribution. In this scenario, the host is thus largely unaffected by microbial variation, but only by the presence of a microbiome. Additionally, not all microbes (e.g. in purple) will not influence host phenotypes. **B)** The microbiome may shift both the variance and the mean of host phenotypes. The blue distribution is suggestive of when host leverage locally adapted microbiomes to specialized phenotypes. **C)** Alternatively, the microbiome may also expand phenotypic variation, conceptualized in green. Expanding host phenotypic variation through the microbiome may allow hosts to explore novel fitness landscapes. These are conceptualized phenotypic distributions, and more experimental work is necessary to confirm how the microbiome affects host phenotypic distributions.

Integrating microbial genetic variation into host evolutionary processes builds on hologenome theory. The hologenome is defined as an evolutionary unit combining the eukaryotic host and all associated microbes^16,24,25^. Under this concept, evolution operates on this single unit because eukaryotic hosts are never isolated from microbes in the natural world^16,23^. Thus, the evolutionary fate of both hosts and microbes is intertwined. The intertwined fate of a single host-microbiome evolutionary unit is the focus of much criticism of hologenome. Because of complicated vertical and environmental transmission routes of the microbiome, the host and microbiome rarely operate as one selective unit^6,7,9^. Moreover, the differences in evolutionary and ecological time scales between host and microbiome also challenge the utility of hologenome theory^8^. Advances in hologenome theory clarify the role of microbial variation in shaping host evolution: variation in the microbiome is associated with variation in host fitness, and hosts and microbiome evolve in response to selective pressures^14,23,25,26^. In turn, selection may operate on the microbiome, host, and their interaction in different ways^25^, but a phenotypic response is necessary to impact the host evolutionary trajectory^9^.

To formalize these verbal arguments, we propose quantitative genetic models that incorporate the contribution of the microbiome to host phenotype, and subsequent response to selection. Furthermore, we explore how fidelity of microbiome inheritance influences the response to selection. Together, this allows us to make predictions about how the microbiome influences host evolution (Box 1). Our simulations show that the microbiome can modulate the host evolutionary response, but this will depend on both the microbial contribution to host phenotypic variation and complexities of microbial inheritance.

## A Complex Inheritance

For the microbiome to influence host evolution, it must have host phenotypic effects and be inherited. From a quantitative genetics perspective, microbiome heritability is the resemblance of microbiomes between parents and offspring. The microbiome can be shared via strict vertical transmission through embryos, but many microbes are also transmitted through quasi-vertical and environmental modes, like vaginal birth in mammals, regurgitation in birds, environmental inoculation in insects, or coprophagy in many taxa^27,28^. Additionally, host genetics can structure microbial variation, just like other complex traits^29^. Host genetic variation may lead to the preferential association of specific environmentally-acquired microbes^30^, leading to high parent-offspring similarity. Indeed, in hosts dominated by environmentally acquired microbes, the contribution of host genetic variation to the relative abundance of particular microbes is as high as 42% in humans^31,32^, 39% in *Drosophila^33^*, and 25% in maize^34^. Interestingly, these and other studies show that not all components of the microbiome are heritable with only a portion of the microbiome faithfully transmitted, with estimates ranging from 8-56% of microbes^32,34–36^.

The inheritance of the microbiome is complex and still poorly understood for several reasons. Some microbes are transmitted exclusively between parent and offspring, like typical host genetic inheritance, while others are environmentally acquired^8,37^. Environmental factors, like diet or climate, also substantially influence the reservoir of possible microbial partners, leading to changes in the microbiome independent of any host evolution^38–40^. Host phenotypes also may be more strongly shaped by distinct microbial variation early in life, but because microbial dynamics operate at shorter timescales than host generations, those relevant microbes may not be faithfully transmitted across host generations^8,41,42^. Such complexities muddle the inheritance of the microbiome, challenging how frequently the microbiome could influence host evolution^6–9,25^. Our simulations show how variation in microbial inheritance modulates the microbial contribution to host evolution (Box 1).

Microbes with beneficial phenotypic effects should be faithfully inherited. However, the drivers of this relationship between inheritance and phenotypic effects is not always clear. For example, in a UK twin study investigating the influence of host genetics on the microbiome, *Methanobrevibacter* species were identified to have the highest heritability and strong association with low body-mass index^31,32^. Similarly, though another study in humans found little influence of host genetics on microbiome composition, microbial variation still explained 22-36% of metabolic traits^39^. Microbial variation explained 33% of weight gain in pigs^35^ and 13% of methane emissions in cows^36^ but also occurred largely independently of host genetic control of the microbiome. In other words, we still do not know whether the most heritable microbes explain the most significant variation in host traits. However, despite the complicated inheritance of the microbiome, these studies suggest that for a range of host traits, the microbiome contributes almost as much to phenotypic variation as host genetics. Our simulations also suggest that when the microbiome shapes host phenotypic variation and is faithfully inherited, the microbiome can contribute substantially to the host selection response (Box 1), but there is currently little empirical data to validate these conclusions.

## Measuring the Microbiome Response To Selection

With recent advances in sequencing technologies, microbial variation can be quantified at many different scales, from strains to whole community composition to complete metagenomes (see^43^ for a recent review). A major challenge is to identify which scale of microbial variation (i.e. strain, community, metagenome) influences host phenotypes^8,43,44^.

#### BOX 1: QUANTITATIVE GENETICS OF HOST-MICROBIOME INTERACTIONS

Quantitative genetic models traditionally decompose host phenotypic variance into genetic (V_G_) and environmental (V_E_) components. However, this simple model generally does not consider the contribution of the microbiome to host phenotypic variation. Extending this quantitative genetic framework to include the microbiome, we use simulations to explore the microbial contribution to host response to selection (see Supp. Methods for more detail).

Ignoring the contribution of epistasis and dominance effects and assuming no covariances between host genetic, microbial, and environmental effects, the total phenotypic variance can be modeled as:

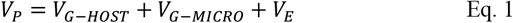

Where V_G-HOST_ is the genetic variance in the host contributing to phenotype, V_G-MICRO_ is genetic variance in the microbiome contributing to phenotype, and V_E_ is the environmental variance. V_G-MICRO_ is frequently quantified as relative abundance of specific microbes. However, with this framework, V_G-MICRO_ can be extended to genetic variation in the microbiome. If we start by assuming all microbial genetic variance is transmitted from parent to offspring, host heritability can be calculated as:

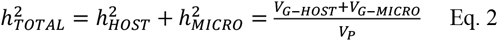

When the microbiome is transmitted across generations, so is the genetic variation the microbiome harbors. Increasing the microbial contribution to phenotype (V_G-MICRO_) increases total host heritability (**Fig. A**)

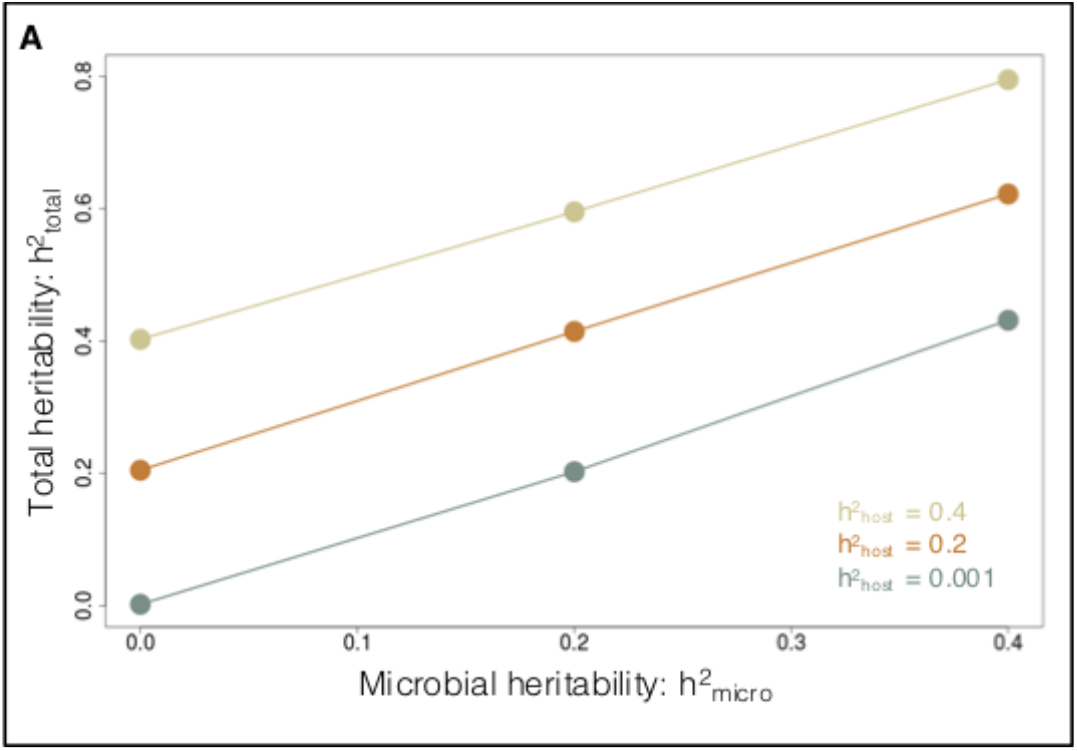

Given the complicated inheritance of the microbiome, we further explored how variation in transmission affects short-term host response to selection. The breeder’s equation can be adjusted to reflect the variation in microbiome transmission:

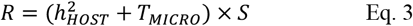

Where R is the response to selection, S is selection coefficient, h^2^_HOST_ is the portion contributed by host genetic variation, and T_MICRO_ is a function modulating the contribution of the microbiome:

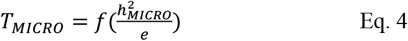

T_MICRO_ scales h^2^_MICRO_, the microbial contribution to host heritability, by *e* microbial transmission noise. In our simulations, we varied transmission noise across two conditions: high and negligible h^2^_HOST_ (high =0.4, negligible=0.0001), while keeping h^2^_MICRO_ high (0.4).

Not surprisingly, the contribution of h^2^_MICRO_ to total heritability decreases as transmission noise increases. The microbiome can modulate the host response to selection, visualized in **B** and **C**. For traits with high host genetic contribution (h^2^_HOST_ = 0.4), the microbiome, if faithfully transmitted, contributes to the response to selection (**Fig. B**). However, as transmission noise increases, the microbial contribution decreases and modulates the response to selection. Second, for traits with negligible h^2^_HOST_, the microbiome alone has the potential to contribute significantly to selection response. The contribution will depend on transmission noise (**Fig. C**). Especially when h^2^_HOST_ is low, the host response to selection will depend on how faithfully the microbiome is transmitted across host generations.

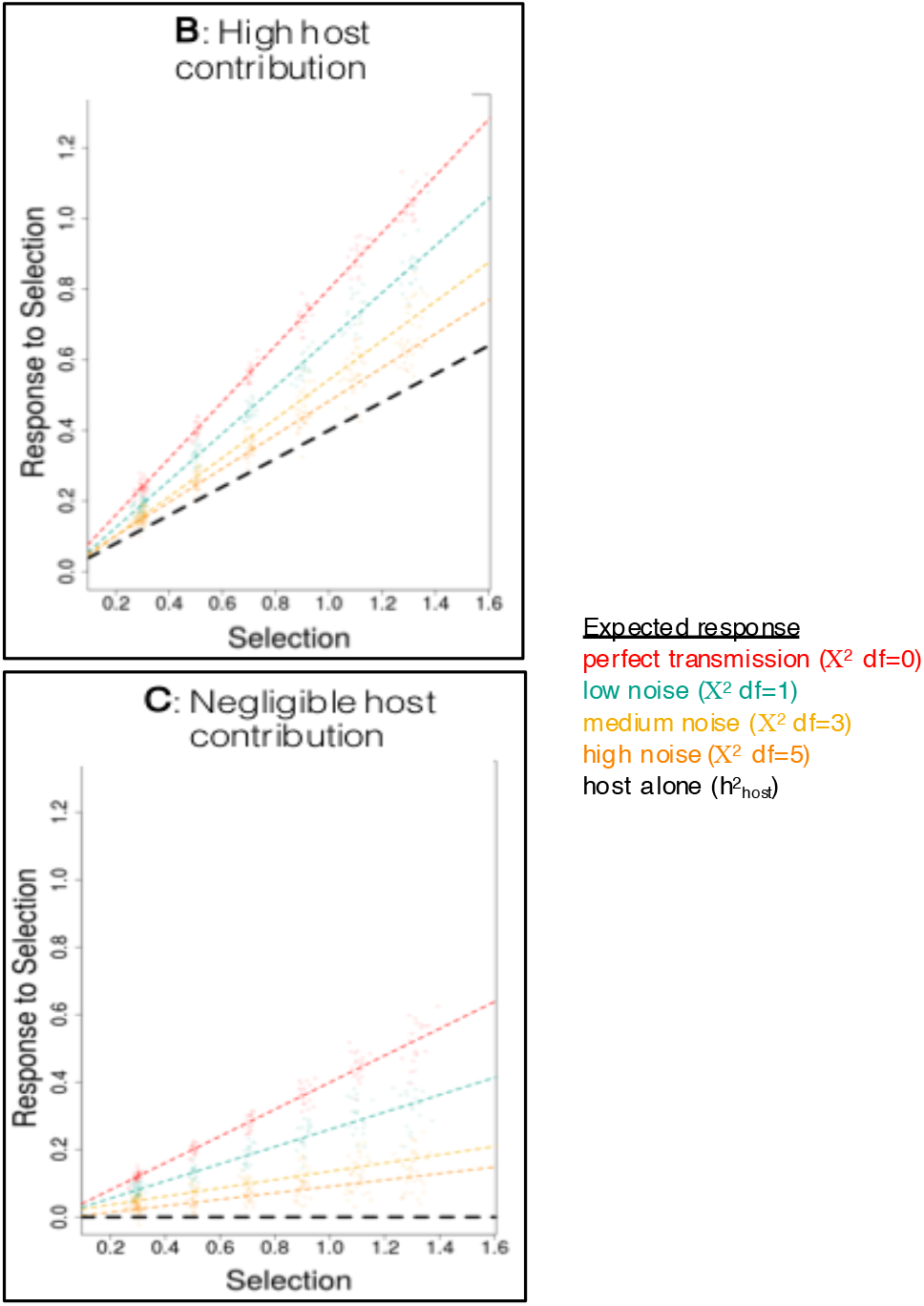

Our simulations highlight the importance of incorporating the microbiome into host evolutionary responses. To contribute, the microbiome must have phenotypic effects and be faithfully transmitted. Although there is a growing interest in quantifying the heritability of the microbiome, there is currently little empirical data available to estimate the variation in parameters used in these simulations. Future work is necessary to better understand the complex relationship between host genetic variation, microbial variation, and heritability.

The majority of host-microbiome interactions are characterized through marker gene studies (i.e. 16S rRNA for bacteria, ITS for fungi). Marker genes can classify microbial communities at broad taxonomic levels, are relatively inexpensive to sequence, and have established bioinformatic and analytical pipelines^43^. This approach demonstrates that microbial communities respond to many different kinds of selective pressures, like drought^45,46^, antibiotics^47,48^, different diets^49^, or warming environments^50^. Some studies show that microbial community diversity decreases in response to stressful environments^46–48^, while others show only shifts in community composition^45,49,50^. Marker gene studies are well-suited for taxonomically defining communities (though see^51^), but changes in microbial community composition do not always correlate to changes in host phenotype. For example, as measured by 16S rRNA marker genes, the macroalgae *Ulva* is colonized by different bacteria in different environments, but metagenomic data shows the same core functional genes encoded by different bacterial taxa^52^. Similar patterns are observed in human microbiomes^53^ and the tree phyllosphere^54,55^. If the microbial genetic repertoire or host phenotypes do not respond to corresponding changes in the microbial communities, then the microbial variation likely has little adaptive value for hosts. However, this conclusion may be strongly impacted by how microbial variation is characterized. Therefore, techniques beyond marker gene classification are needed to determine if and how microbial variation influences the host response to selection.

Strain-level analyses of microbial variation (i.e. polymorphisms and other genetic variants beyond marker genes) may provide more functional insight into the microbiome response to selection. For example, strain-level variation in honey bee microbes influences the response to pathogens or different diets for the hosts^56^. Strain-level analyses also provide insights into the stability of the microbiome across the lifetime of the host^57–59^ as well transmission dynamics^60,61^. Metagenomes, which fully characterize the extended genotype, will provide crucial insights into functional variation of the microbiome^43,56,62,63^. For example, metagenomics demonstrated how bacterial metabolism varies across different niches within the same host^64^. This approach however remains costly, computationally intensive, and the results can be difficult to interpret. The emerging field of metatranscriptomics aims to study gene expression in the complex microbial communities^43,65,66^. Linking the metagenome to the metatranscriptome will be particularly insightful to understand how microbial variation influence host phenotypic variation^43,67^. For example, by comparing metagenomes to metatranscriptomes in the human gut, it was discovered that only a few specialized bacteria express unique biosynthesis pathways that may reflect the microbiome response to changing diets or other environmental perturbations in hosts^67,68^. Experimental approaches that compare how microbial genetic variation alters host gene expression (interspecies eQTLs^69^), and vice versa, provide a promising way forward to understand the links between host-microbiome response to selection at the whole system scale.

## Linking the Microbiome to Host Adaptive Responses

Several analytical and theoretical gaps limit our current understanding of how the microbiome influences host evolution. First, while remarkable progress has been made in detailing microbial diversity, studies examining genetic variation in the microbiome (i.e. SNPs) beyond marker genes remain very limited^43,59^. The growing number of metagenomic studies will answer some questions, but more advanced bioinformatic tools are needed to connect strain-level microbial variation to host phenotypic effects. On the theoretical side, many aspects of microbial evolution--like demography, pangenomes, horizontal gene transfer, unique population structure--may poorly match the assumptions of more traditional population genetic models of free-living organisms^59,70–72^. Can we, for example, apply indirect selection^13^ or multilevel selection models (i.e. Price equation)^44^ to ask how independent or synergistic are host and microbiome evolution? One novel approach to understand the signatures of microbial variation shaping host evolution is interspecies linkage disequilibrium^73^. Interspecies linkage disequilibrium may occur when beneficial combinations of host and microbiome alleles jointly increase host fitness. If so, selection may operate on the linkage, resulting in nonrandom allelic assortment of beneficial microbes and host genetic variation. The challenge is to identify which and how frequently variation in the microbiome increases host fitness, and whether this increases the probability of transmission across host generations.

#### BOX 2: Testing The Microbiome Influence On Host Evolution

Experimental approaches developed for variance partitioning in quantitative genetics can be a powerful way to assess the influence of the microbiome on host evolution. **A)** Experimental evolution will provide critical insights into how host and microbiome respond to stressful environments. By including the microbiome (visualized as different colored circles) in experimental evolution, then genetic responses in both host and microbiome can be measured following selection. At the end of experimental evolution, we expect both host and microbiome to adapt (visualized as blue fly and blue/purple microbes). To test how the microbiome interacts with host genome to influence host phenotypes, one can perform fully factorial, reciprocal transplants between host, microbiome, and environment. **B)** Key insights will be gained from examining the evolutionary trajectory of alleles that emerge or change in frequency during experimental evolution. **C)** To test how microbial variation influences host phenotype, hosts can be inoculated with different levels of microbial variation. Removing the microbiome through antibiotics (or other manipulations) will provide insights into how hosts respond to perturbation to their microbiomes. **D)** Finally, we advocate that diallel crosses can be used to show how host and microbial genetic variation interact. Diallel crosses are performed by crossing all possible combinations between inbred lines to each other in a common environment (represented by fly colors). Rearing F1s in different microbial environments will enable partitioning of the additive and nonlinear, epistatic components between host and microbial genetic variation.

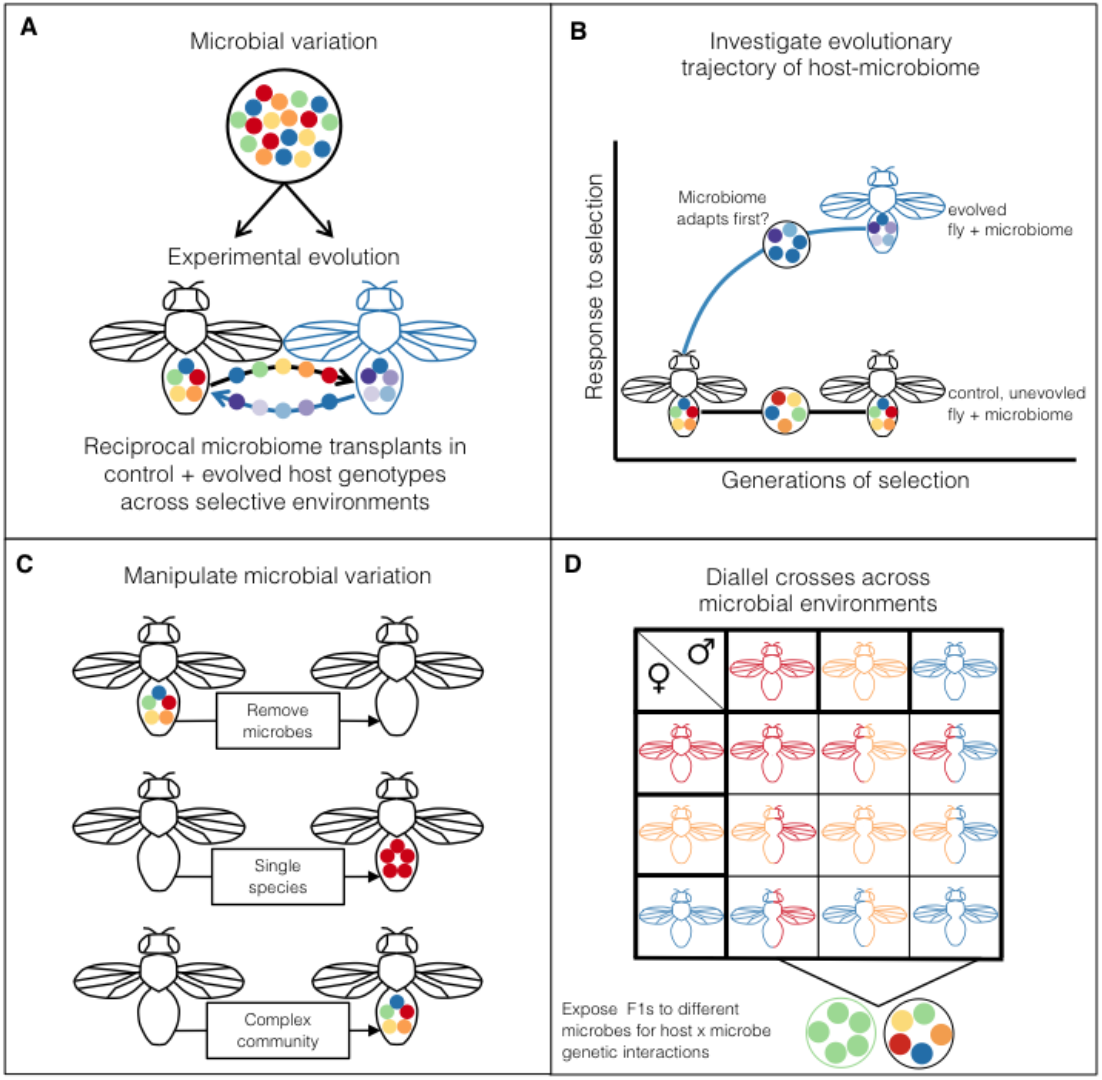

Developing meaningful null models to describe host-microbiome evolution is a nontrivial challenge. First, we lack empirical data to calibrate a null model, like when the microbiome does not influence host phenotypes. Second, the null model should consider the balance in evolutionary interests between host and microbiome. Hosts and microbiomes will rarely have perfectly aligned evolutionary interests, suggesting a balance between cooperation and conflict^6,8,41^. For example, intergenomic conflict is likely to have important consequences for host-microbe coevolution because environmentally acquired microbes face vastly different evolutionary pressures inside and outside of hosts. Intergenomic conflict may maintain variation in microbial associations, but how often this influences host-microbiome evolution is not well understood^74^. Ultimately, the null hypothesis should consider the balance between cooperation, conflict, and indifference between the host and microbiome during adaptation.

Ultimately, rigorous sampling across time and space will be key to illuminating drivers of host-microbiome evolution. Within the last 10 years, researchers have started to identify how the microbiome influences host evolutionary processes in diverse taxa and environments. We synthesized the literature and find evidence suggestive of two common scenarios in how the microbiome may shape host evolution: 1) hosts leverage locally adapted microbes, and 2) the microbiome increases host intraspecific phenotypic variation. These scenarios are not mutually exclusive, but as we explain below, suggest different evolutionary trajectories for hosts.

## Hosts Leverage Locally Adapted Microbes

In stressful environments, hosts may leverage locally adapted microbes. Microbes with larger effective population sizes, rapid generation time, and pangenomes may evolve novel functions faster than their hosts^75–78^. If hosts can capitalize on locally adapted microbes, then hosts can increase survivorship and rapidly adapt to novel environments.

There are a number of remarkable studies that illustrate the substantial impact of locally adapted microbes on hosts, suggesting that microbes may underlie host adaptation to populations in stressful environments. For example, bean bugs can gain pesticide resistance through acquiring a pesticide degrading soil bacterium, *Burkholderia^79–81^*. Many other hosts utilize their microbiome to detoxify harmful chemicals. The woodrat microbiome degrades terpenes, enabling the woodrat to live in a specialized dietary niche^82–86^. Bark beetles^87^, pine weevils^88^, and coffee berry borers^89^ also use their microbiomes to detoxify plant secondary compounds in specialized niches. The microbiome can also facilitate survival in other kinds of stressful environments. Plants on geothermal soils are associated with the thermotolerant endophyte *Curvularia^90^*. Thermotolerant *Curvularia* can increase survival up to 40°C in non-adapted tomatoes, while *Curvularia* isolated from non-geothermal soils did not^91^. Additionally, salt tolerant fungi can also confer salt tolerance to non-adapted plants^91^. Coral modify their microbiomes in response to temperature stressors and pathogens, facilitating survival in rapidly changing oceans^92–94^. These examples show that hosts can utilize specific microbial genotypes or communities with large phenotypic effects to specialize and persist in novel niches.

Relying on locally-adapted microbes may facilitate evolution in the short term--but what are the evolutionary consequences of host reliance on locally adaptive, environmentally-acquired microbes? The evolutionary benefit may allow hosts to rapidly cross the fitness landscape. Then, hosts should evolve to maintain the locally adapted microbes and their modification to the environment, much like genetic accommodation or niche construction^95^. However, relying on environmentally acquired microbes increases stochasticity in the microbiome, exposing offspring to pathogens and non-adaptive microbes. If specific microbes are beneficial, hosts should evolve mechanisms to ensure faithful microbial transmission, like traditional symbioses^96^. While still contentious if most microbiomes behave like traditional symbioses, ultimately, strict vertical transmission and extreme reliance on specialized microbes is likely an evolutionary dead end, constraining hosts and microbes to the ecological conditions that generated the symbiosis^97^. Overall, the challenge remains in determining how microbial genetic variation is linked with and influences host genetic change during adaptation.

## Microbial Variation Shapes Host Intraspecific Phenotypic Variation

In the case of locally-adaptive microbiomes, specific microbes enable host survival by optimizing phenotype to match specific stressors, shrinking host phenotypic variation (Fig. 1). However, in other cases, stochastic variation and priority effects during microbial community assembly may create microbial variation between hosts^8,42,98^. Such an increase in microbial variation may increase phenotypic variation in host population, enabling more individuals to explore broader phenotypic spaces^99,100^ (Fig. 1). Microbial variation would then enable the exploration of new fitness landscapes where hosts may more rapidly find novel solutions to stressful environments.

Microbial variation increases host phenotypic variation in many organisms. For example, in *Drosophila*, different microbial communities increase the variance in larval development time, pupal weight, and adult weight compared to sterile flies^101,102^. In *Daphnia*, different bacteria increase variance in body size and hatching success compared to sterile treatments^103^. Microbiomes evolved for rapid flowering time in *Arabidopsis* increased variance in biomass compared to microbiomes evolved for slow flowering time^104^. In zebrafish, different microbes are associated with varying immune responses^105^. In all these cases, hosts are permissive to colonization by many microbes, and in turn, phenotypes respond to microbial variation. A major question for host-microbiome evolution is why some host phenotypes are responsive to microbial variation while others maintain more constant traits (i.e they are more canalized)?

Maintaining responsive phenotypes to microbial variation may allow hosts to better match phenotypes to current environmental stressors^17,18^. For hosts where offspring might occupy different environments than parents, microbes may provide critical cues for local environments. For example, microbes evolved under drought led to early flowering in both *Arabidopsis* and *Brassica^104,106^*, suggesting that microbes signal local growing conditions and changes in life-history strategies. For *Daphnia*, microbes acquired from their local environment increased fecundity compared to those acquired via maternal transmission before diapause^103^. To best match phenotype to local environment, hosts may use microbial variation to alter developmental timing. Development in many taxa, from insects^107–113^ to plants^114–116^ to crustaceans^117,118^, changes in response to microbial variation. Developmental plasticity, through microbial variation, may expose novel phenotypic variation^119^, and this may in turn generate phenotypes that allow rapid acclimation to variable environments. Hosts may evolve to maintain variable microbial associations that best match phenotypes to current environments.

Phenotypes that are responsive to diverse microbial associations may alternatively pose a risk to the host. Microbial variation may lead to mismatches between host and environment that reduce host fitness, like in the case of westernized human microbiomes leading to increased risk of obesity in immigrants in the United States^120^. Ultimately, relying on environmentally acquired microbes depends on the microbes providing honest signals of local environmental conditions^11,119,121^. Identifying how hosts utilize environmentally acquired microbes to match phenotypes to environments is understudied, but essential to understanding the evolutionary benefits of the microbiome.

## Using Experimental Evolution To Characterize Microbial Influence

Above, we identified common scenarios of how the microbiome may influence host evolutionary processes. To provide more mechanistic insight into host-microbiome evolutionary processes, experimental evolution can provide a powerful approach to understand how microbial change interacts with host evolution^14,113,122^. However, few studies have actually explicitly evaluated evolutionary responses in the microbiome. To illustrate how the microbiome responds to experimental evolution, we re-analyzed genomic data from 10 Evolve and Resequence (E&R)^123^ experiments in *D. melanogaster^124–132^* to characterize variation in the microbiome. We compared bacterial diversity between control and evolved populations to measure the microbial response to selection (see Supp. Methods for more detail).

We found that bacterial diversity frequently responded to experimental evolution (Fig. 2A). Evolved populations often exhibited reduced bacterial diversity (4/10 studies), though in one case (accelerated development time), increased bacterial diversity. Because each E&R experiment varied in length of selection (from 5-605 *Drosophila* generations), we also tested if change in microbial diversity was correlated with host generations of selection. Change in microbial diversity was only weakly correlated with host generations of selection (Fig. 2B, *r*=0.26, p=0.02). The specific nature of the selective pressure is more important in driving changes in the evolving microbiome. For example, the evolved microbiome in starvation resistance exhibited the greatest change in bacterial diversity (Fig. 2B), and the *Drosophila* microbiome is tightly linked to the regulation of metabolic networks^133^. For other traits, like egg size, the microbiome did not respond to experimental evolution. This meta-analysis suggests that the effect of selection on microbiome is likely trait specific.

**FIGURE 2:**
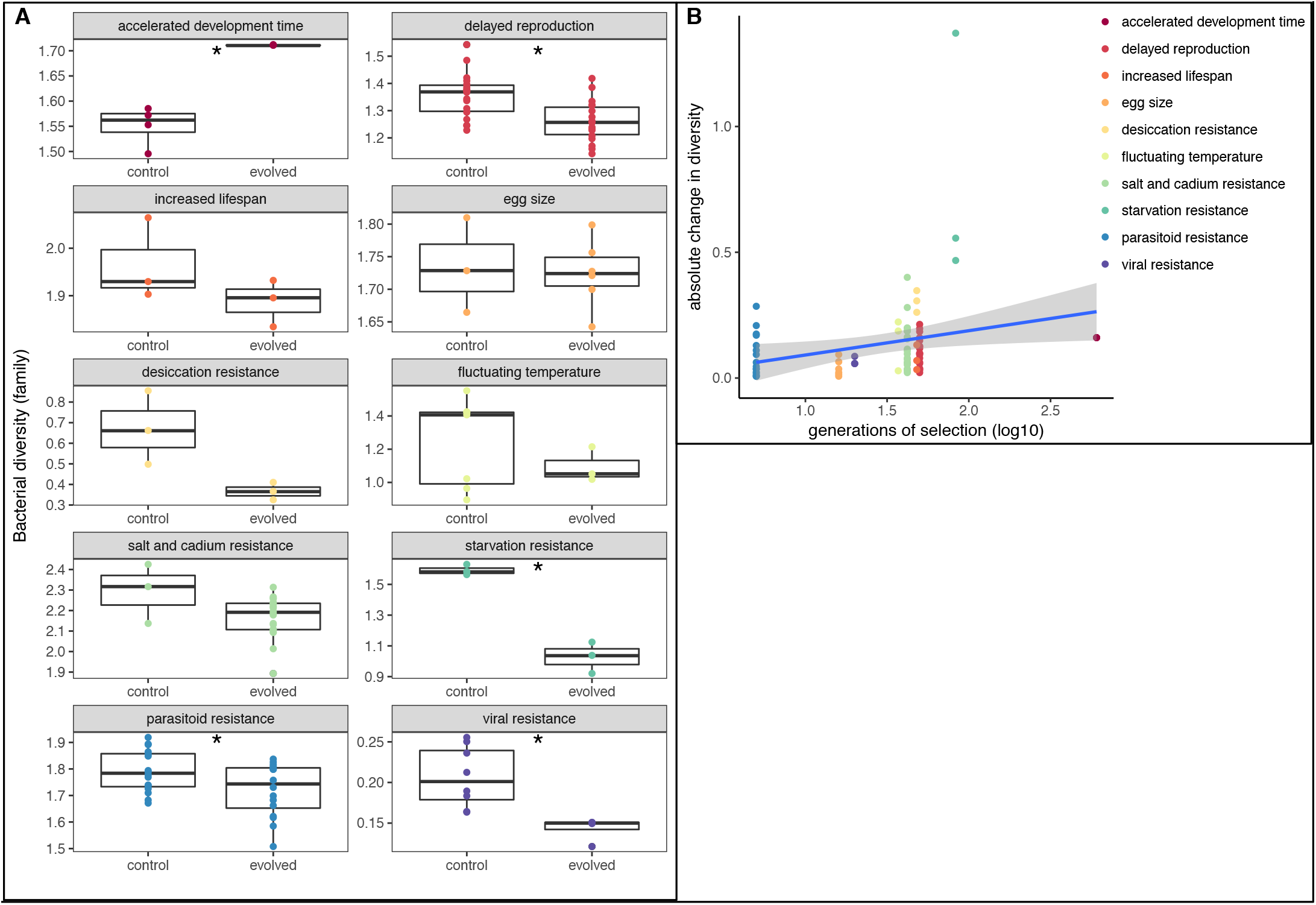
We reanalyzed 10 E&R experiments to determine how bacterial diversity responded to experimental evolution in D. melanogaster. A) 5/10 experiments had significantly different bacterial diversity between control and evolved lines (denoted with asterisk). See Supp. Table 2 for summary of statistical tests. B) Evolved microbial diversity was only weakly correlated with host generations of selection (r = 0.24, p = 0.02).

Our meta-analysis suggests that the microbiome evolves with the host during experimental evolution. However, like other studies, we are only observing the endpoint of the host-microbiome evolutionary trajectory. Ultimately, understanding how microbiome alters the evolutionary trajectory in hosts is the critical missing link^122,134–141^. Monitoring the temporal dynamics during experimental evolution can provide the missing link--specifically, does the microbiome adapt faster than the host? And how frequently does the host utilize this rapid microbial evolution? A recent study, where the temporal evolutionary dynamics were monitored, suggests that *Lactobacillus* bacteria in *Drosophila* evolve rapidly to nutrient poor diets, and *Drosophila* can leverage rapid microbial adaptation independent of their own evolution^113^. More experiments are needed that combine genotyping and phenotyping host-microbiome interactions to understand the temporal evolutionary dynamics. We outline how experimental evolution, combined with other tools commonly used in quantitative genetics, can provide deep insights into how the microbiome influences the host evolutionary trajectory (Box 2).

## A Practical Outlook

Based on the examples discussed above, we show that the microbiome can influence host phenotypes and their evolutionary trajectory across taxa and diverse environments. Natural variation in the microbiome may also provide new tools to combat a range of challenges with practical applications. Indeed, there are several promising avenues of research leveraging natural microbial variation in fields ranging from conservation biology to agriculture. For example, variants of *Pseudomonas* bacteria isolated from caves help protect vulnerable bats from the devastating fungal infections of white-nose syndrome^142,143^. A similar strategy is also in development to protect amphibians from chytrid pathogens^144^. From a public health perspective, manipulating the microbiome in disease vectors may reduce vector competence^145^. The success of cancer management can be influenced by the presence of particular gut microbes in humans, and finding novel probiotics to improve the efficacy of chemotherapy is under development^146^. Isolating host-associated microbiomes from organisms in extreme environments, like geothermal soils^90,91^, will serve as reservoir of adaptive microbes for agriculture in changing environments^14,147^. Natural variation in the microbiome represents a largely untapped resource with many novel solutions to address applied challenges.

## Conclusions

Here, we have explored how the microbiome influences host evolution. Through synthesizing the literature, modeling, and metaanalyses, we identify conditions when the microbiome has the greatest potential to influence host evolution. Nevertheless, several key evolutionary questions remain to be addressed (Box 3): How do ecological interactions between microbes shape host phenotypes? What is the genetic basis of host responsiveness to microbial variation? How does the microbiome change host adaptive response? And, do the results of lab studies apply to microbiomes in the wild? Answering these questions will require a diversity of approaches using tools from quantitative genetics, community ecology, and genomics. We advocate for integrative experiments that leverage experimental evolution and natural variation to understand how the microbiome shapes host evolution (Box 2). More work, particularly in natural systems, is especially critical to find generalities of host-microbiome evolution.

In conclusion, the complex interplay between host and microbial genetic variation is surprisingly understudied. We find that the microbiome frequently shapes host phenotypic variation across diverse taxa and environments. In doing so, the microbiome may enable hosts to more rapidly explore the fitness landscape than the host could on its own. Studying the interplay between host and microbial genetic variation is complicated, but continuing advances in sequencing technology will facilitate the necessary characterization. Incorporating microbial variation will provide fundamental insights into how phenotypic variation is generated, and subsequently, how selection operates across ecological and evolutionary scales.

#### BOX 3: Key questions and research priorities

##### How does the microbiome change the host’s adaptive response to stressful environments?

Our E&R meta-analysis suggests the microbiome also responds to selection. When the microbiome responds to selection, then does the microbiome reach a new adaptive peak before the host? Does the microbiome reinforce or dilute the selective forces acting on the host? Sequencing and microbiome transplantation along the course of experimental evolution will provide key insights into the role of microbiome in host adaptation. Equally important is investigating systems where microbes do not impact host adaptation^6,44,148^.

##### How does microbial community assembly interact with host development to shape host phenotypes?

How do microbe-microbe interactions influence host phenotype? How does community assembly interact with host developmental to shape host phenotype? Integrative work where permutations of bacterial communities colonize controlled host genotypes in Arabidopsis^149,150^ and Drosophila^107,151,152^ are important first steps; the challenge remains in applying this approach across different stressors as well as to non-model systems in the laboratory and field.

##### What is the genetic basis underlying host responses to microbial variation?

Are there microbes (or genes) with disproportionately large effects on host phenotype? Do few host genes enable specificity or promiscuity in microbial associations? What host genes integrate and transform microbial variation into phenotypic variation? To understand how host and microbial genomes interact, we will need to combine common marker gene sequencing approaches with new approaches in characterizing microbial genetic variation at the strain level^153–155^. Computational approaches that ask how variation in the microbiome interacts with host genetic variation, like interspecies linkage disequilibrium^156^ -and interspecies eQTLs^69^, will illustrate mechanistic processes underlying microbiome-driven host adaptation.

##### What ecological and evolutionary forces structure microbiomes in the wild?

What abiotic and biotic pressures influence microbial variation? How faithfully are microbiomes transmitted across generations and/or environments? How frequently do microbiomes influence host fitness in the wild? To answer these questions, researchers should following best practices to sample the microbiome in both hosts and the environment^43,157,158^. Identifying how the microbiome is transmitted and maintained in the wild will be crucial to understanding how the environmentally acquired microbiome shapes evolutionary processes.

## Acknowledgements

We would like to thank Sarah Kocher, Amanda Lea, Jess Metcalf, Lindy McBride, and Luisa Pallares for helpful feedback.

## Code Availability

Code for simulations and E&R microbiome analyses can be found at https://github.com/lphenry/extgeno.

## SUPPLEMENTAL METHODS FOR QUANTITATIVE GENETIC SIMULATIONS

In order to simulate the effect of the microbiome on selection response, we considered families with three offspring each. This scenario is described by a block diagonal kinship matrix 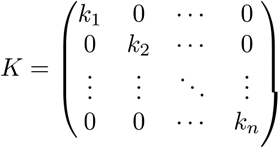, where each block 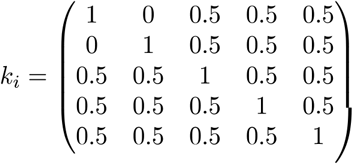 corresponds to one family.

The contribution of the host genome to the phenotype is given by 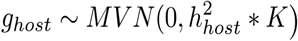, in accordance with quantitative genetics theory^11^. Extending the conventional theoretical framework, we define the phenotypic contribution of the microbiome as 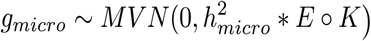, where 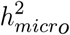 is the phenotypic variance explained by the microbiome in the parental generation. 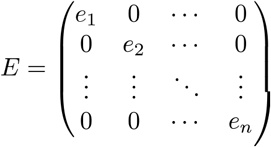 is a block diagonal matrix determining the transmission noise, where each block 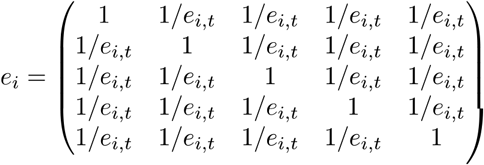 corresponds to one family.

The Hadamard product^22^ *E* o *K* thus produces a matrix where the co-variances are down weighted by a factor ^*e_i_*^,*t* per family. For simplicity, the transmission noise ^*e_i_*^,*t* is sampled from a chi-square distribution, the degrees of freedom determining the amount of noise (df = 1, 3, and 5). Finally, the phenotype is defined as *y* = *g_host_* + *g_micro_* + *e*, where 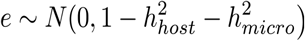, thus keeping total VP is kept constant.

## SUPPLEMENTAL METHODS FOR E&R ANALYSES

We downloaded SRA files for the 10 Evolve and Resequence (E&R) experiments from NCBI. The E&R approach sequences pools of individuals from different selection regimes, but each E&R study had different levels of replication, summarized in Supp. Table 1. Our analyses captured a wide range of different selection pressures, from life-history traits to abiotic pressures to pathogen pressures. Raw sequences were cleaned using Trimmomatic^3^ to remove sequencing adapters, remove low quality reads (average quality per base > 15), and drop reads shorter than 20 bases long. Then, bacterial reads were called using Kraken^4^ to family level. Relative abundance of bacterial families were determined using Bracken^5^. Because the E&R sequencing was not performed explicitly for the microbiome, we needed to remove potential contaminants. We removed any low abundance bacterial family that was assigned fewer than 100 reads. Microbial diversity was calculated using the Shannon diversity index.

To test if microbial diversity was different between control and evolved lines, we compared Shannon diversity. We determined significance using a t-test (see Supp. Table 2 for significance summary). We then tested if two factors explained differences in microbial diversity between control and selected lines: length of selection and *Wolbachia*.

First, length of selection ranged from 5-605 generations. We reasoned that selection response in the microbiome might be influenced by length of selection; the longer the selection, the more divergent the microbiome between control and evolved lines. To test if length of selection was correlated with changes in microbial diversity, we calculated the average microbial diversity for the control lines. We then subtracted the diversity of each evolved line from the averaged control diversity to calculate change in diversity. Because we had positive and negative changes in diversity, we used the absolute difference. We performed a linear regression between change in diversity and the log10 length of selection in R.

Second, we tested the effects of *Wolbachia* on changes in the selection response of the microbiome. As mentioned in the full text, *Wolbachia* is a facultative, intracellular bacteria transmitted exclusively from mother to offspring. *Wolbachia* has no known environmental reservoir and cannot exist outside of the host. For the extended genotype, these intracellular, maternally transmitted bacteria have the same evolutionary trajectory as the host genome; the shared transmission mode limits their ability to rapidly respond to stressful environments. So, we tested if *Wolbachia* influenced the response to selection in the microbiome. To test the role of *Wolbachia*, we removed *Wolbachia* reads from the communities then recalculated Shannon diversity. We compared if bacterial communities without *Wolbachia* significantly differed between control and evolved lines using a t-test.

---

**Supp. Table 1:**
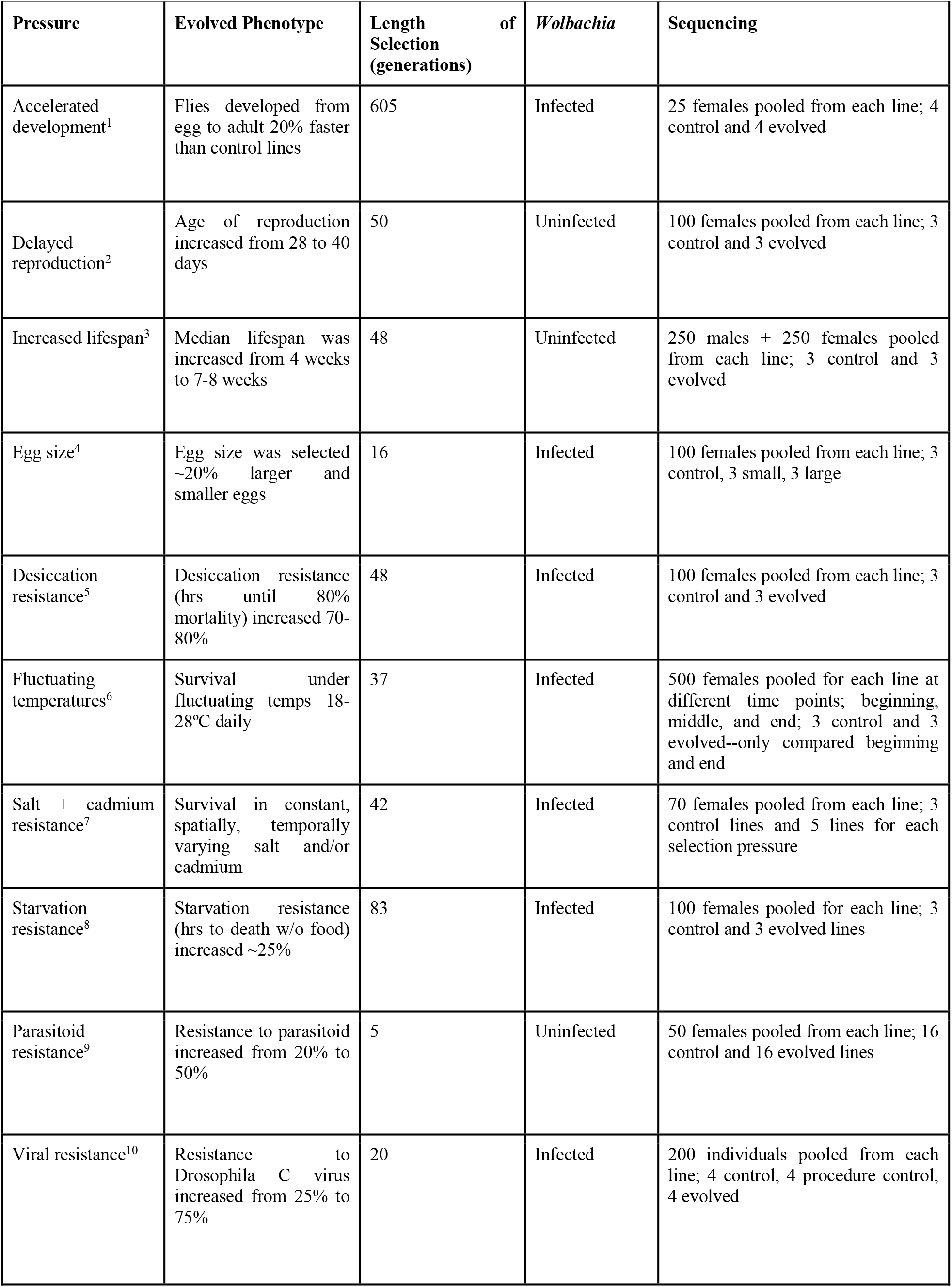
Evolve & Resequence data.

| **Pressure** | **Evolved Phenotype** | **Length of Selection (generations)** | ***Wolbachia*** | **Sequencing** |
| --- | --- | --- | --- | --- |
| Accelerated development<sup>1</sup> | Flies developed from egg to adult 20% faster than control lines | 605 | Infected | 25 females pooled from each line; 4 control and 4 evolved |
| Delayed reproduction<sup>2</sup> | Age of reproduction increased from 28 to 40 days | 50 | Uninfected | 100 females pooled from each line; 3 control and 3 evolved |
| Increased lifespan<sup>3</sup> | Median lifespan was increased from 4 weeks to 7-8 weeks | 48 | Uninfected | 250 males + 250 females pooled from each line; 3 control and 3 evolved |
| Egg size<sup>4</sup> | Egg size was selected ~20% larger and smaller eggs | 16 | Infected | 100 females pooled from each line; 3 control, 3 small, 3 large |
| Desiccation resistance<sup>5</sup> | Desiccation resistance (hrs until 80% mortality) increased 70-80% | 48 | Infected | 100 females pooled from each line; 3 control and 3 evolved |
| Fluctuating temperatures<sup>6</sup> | Survival under fluctuating temps 18-28°C daily | 37 | Infected | 500 females pooled for each line at different time points; beginning, middle, and end; 3 control and 3 evolved--only compared beginning and end |
| Salt + cadmium resistance<sup>7</sup> | Survival in constant, spatially, temporally varying salt and/or cadmium | 42 | Infected | 70 females pooled from each line; 3 control lines and 5 lines for each selection pressure |
| Starvation resistance<sup>8</sup> | Starvation resistance (hrs to death w/o food) increased ~25% | 83 | Infected | 100 females pooled for each line; 3 control and 3 evolved lines |
| Parasitoid resistance<sup>9</sup> | Resistance to parasitoid increased from 20% to 50% | 5 | Uninfected | 50 females pooled from each line; 16 control and 16 evolved lines |
| Viral resistance<sup>10</sup> | Resistance to Drosophila C virus increased from 25% to 75% | 20 | Infected | 200 individuals pooled from each line; 4 control, 4 procedure control, 4 evolved |

**Supp. Table 1:**
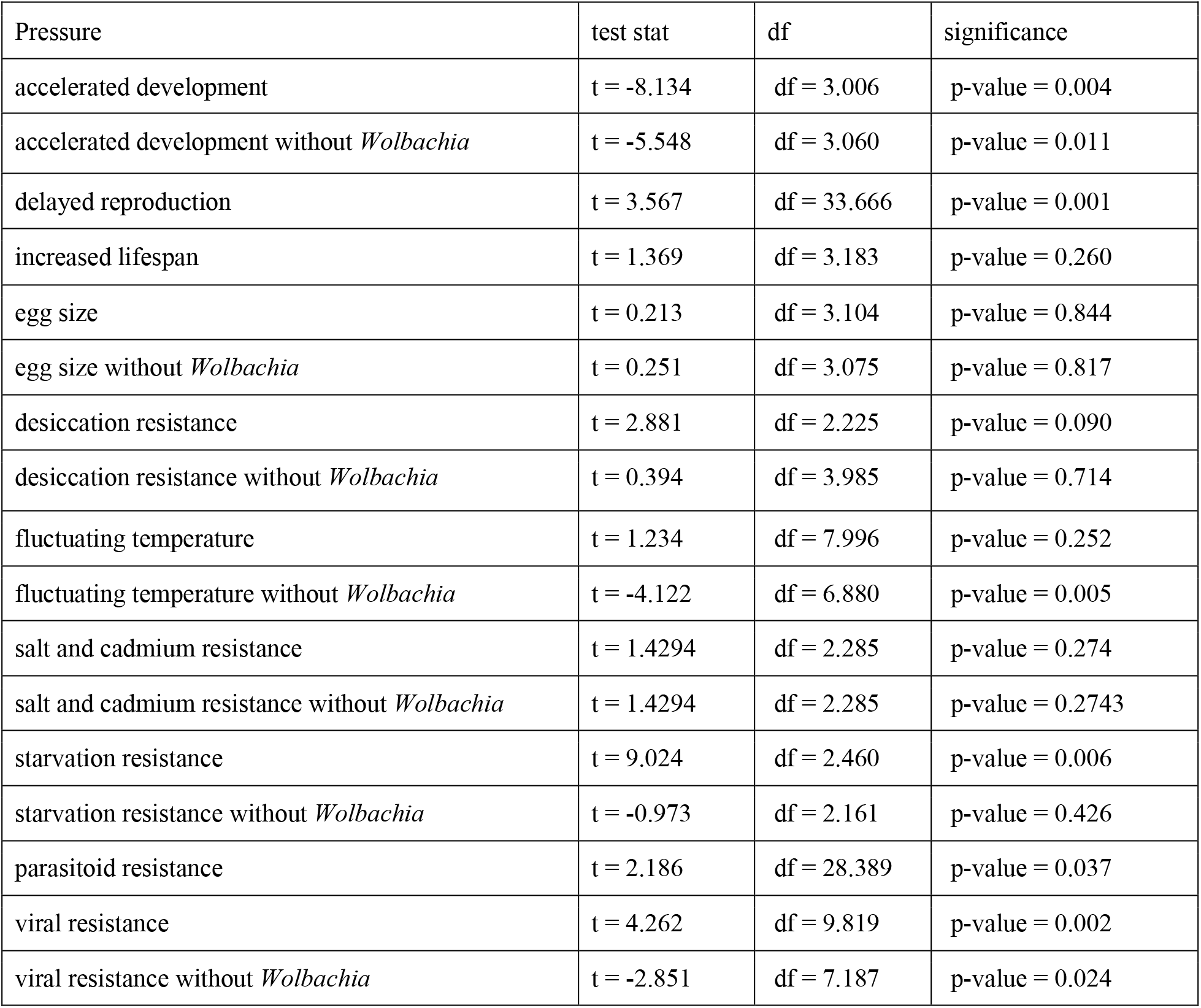
Statistical differences between control and evolved microbiomes.

**Supp Fig. 1:**
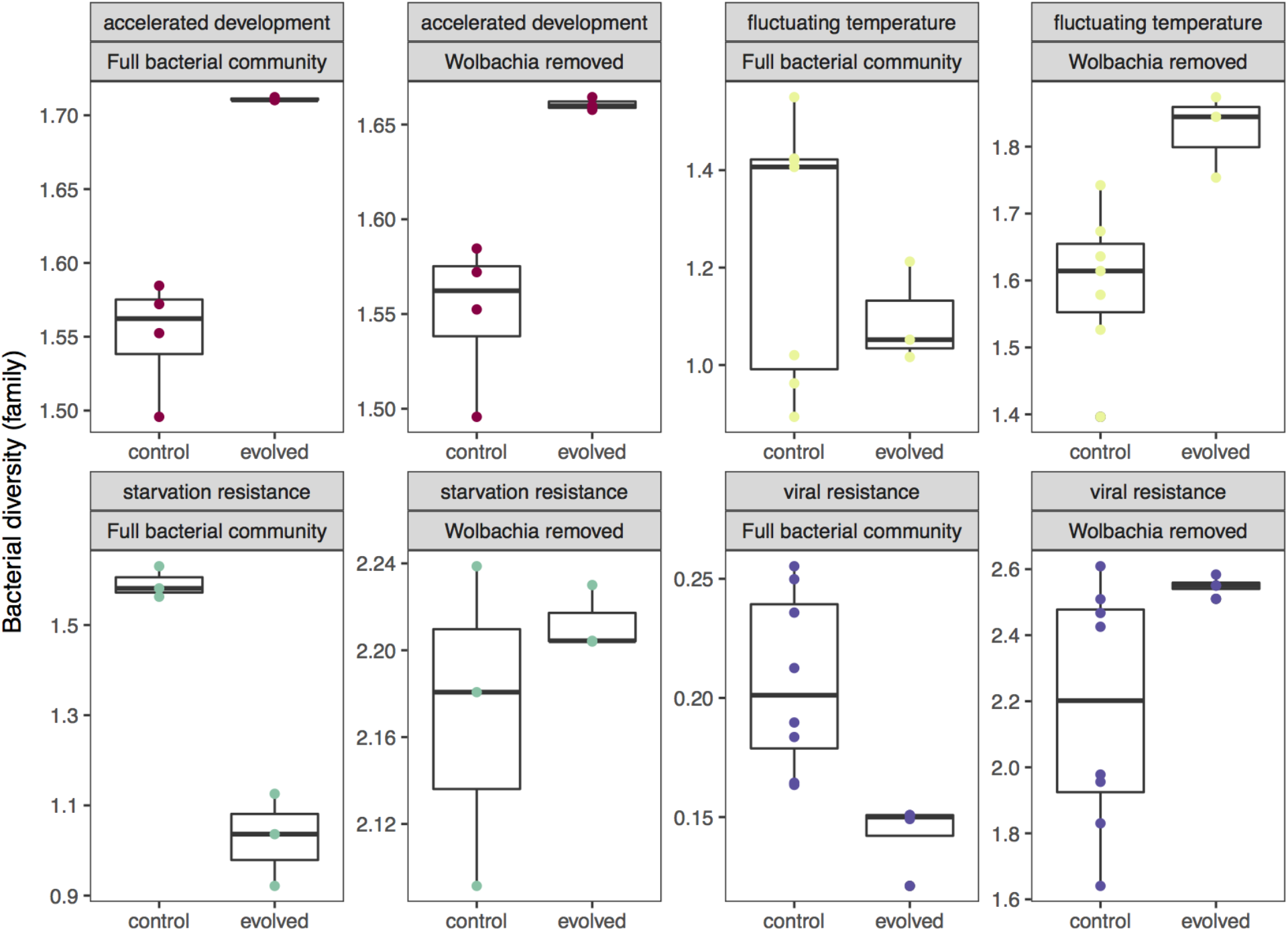
Removing *Wolbachia* frequently changes the response of the evolved microbiome (Supp. Table 2 for statistical summary). First, in accelerated development, removing *Wolbachia* did not affect bacterial diversity between control and evolved lines. Second, removing *Wolbachia* lead to increase in environmentally acquired bacterial diversity in the evolved lines under fluctuating temperatures. Third, removing *Wolbachia* shows that environmentally acquired bacteria did not respond to selection in starvation resistance as there was no different between control and evolved lines. Finally, in viral resistance, including *Wolbachia* showed a reduced microbial diversity in evolved lines. However, removing *Wolbachia* increased environmentally acquired bacterial diversity in evolved lines.

